# Disentangling the process of speciation using machine learning

**DOI:** 10.1101/356345

**Authors:** Megan L. Smith, Bryan C. Carstens

**Affiliations:** Department of Evolution, Ecology, & Organismal Biology, The Ohio State University, 318 W. 12th Avenue, 300 Aronoff Labs, Columbus, OH 43210-1293, USA.

**Keywords:** Speciation, Species Delimitation, Machine Learning

## Abstract

Historically, investigations into the processes driving speciation have largely been isolated from systematic investigations into species limits. Recent advances in sequencing technology have led to a rapid increase in the availability of genomic data, and this, in turn, has led to the introduction of many novel methods for species delimitation. However, these methods have been limited to divergence-only scenarios and have not attempted to evaluate complex modes of speciation, such as those that include gene flow during early stages of divergence (sympatric speciation) or population size changes (founder effect speciation). To address this shortcoming, we introduce *delimitR*, an approach that enables biologists to infer species boundaries and evaluate the demographic processes that may have led to speciation. *delimitR* uses the binned multidimensional Site Frequency Spectrum and a machine-learning algorithm (Random Forests) to compare speciation models. We use simulations to evaluate the accuracy of *delimitR*. When comparing models that include lineage divergence and gene flow for three populations, error rates are near zero with recent divergence times (<100,000 generations) and a modest number of Single Nucleotide Polymorphisms (SNPs; 1,500). When applied to a more complex model set (including divergence, gene flow, and population size changes), error rates are moderate (~0.15 with 10,000 SNPs), and misclassifications are generally between highly similar models. We also evaluate the utility of *delimitR* using three previously published datasets and find results that corroborate previous findings. Our analyses indicate that *delimitR* can serve as an important conceptual bridge uniting various investigations into the process of speciation.

## Significance Statement

For centuries, discovering and describing earth’s biodiversity has been a central goal of biologists. Recent advances have led to an explosion of both molecular data and of molecular-based approaches to species delimitation. Until now, methods for species delimitation have ignored many of the processes considered to be of importance in speciation, including gene flow and population size changes. We introduce a new approach for species delimitation with molecular data that simultaneously infers species boundaries and the processes by which these boundaries arose. Our method has low error rates when applied to typical species delimitation problems and corroborates previous results when applied to published datasets. This method is available as an R-package on github.

## Introduction

Historically, investigations that seek to identify species limits have been largely independent from those that explore the process of speciation. Due to recent advances in Next-Generation Sequencing (NGS) techniques, evolutionary biologists can now collect tens of thousands of SNPs from within and between species at a reasonable cost, which has led to a rapid increase in the number of phylogeographic datasets that may be informative at the level of the species-boundary (1). The field has seen a corresponding increase in available methods for species delimitation that use molecular data (e.g. 2,3, reviewed in 4), with most relying on the multi-species coalescent model (MSCM). While the MSCM is a powerful framework for estimating population sizes and divergence times while accounting for ancestral polymorphism and incomplete lineage sorting (5), it is limited to situations in which gene flow ceases immediately upon population divergence, corresponding to an allopatric mode of speciation. Allopatric speciation may be common, but it is certainly not the only mechanism by which species arise in nature (6–8). Processes other than lineage divergence play an important role in some modes of speciation. For example, gene flow during divergence is a hallmark of sympatric speciation (6), and has been implicated in several empirical systems, including *Myotis* bats (9), Tennessee cave salamanders (10), flowering plants on Lord Howe Island (11), and *Heliconius* butterflies (12). Gene flow may also play an important role during the later stages of divergence via reinforcement, the process by which gene flow between divergent populations results in hybrids with lower fitness and thereby increases positive assortative mating among members of the same lineage, leading eventually to reproductive isolation between cooccurring lineages (e.g. 13, 14). Other population-level evolutionary processes, including population size changes, are also important in some proposed models of speciation. For example, in founder effect speciation (15, 16), a small number of individuals colonize a new area (generally via long-distance dispersal), and the new population is immediately isolated from the ancestral population (15, 17). Genetic drift and natural selection lead to a shift in adaptive peak, and speciation occurs (16). Ignoring these processes while considering only lineage divergence represents the major barrier to uniting systematic investigations into species limits and evolutionary investigations into the process of speciation.

Additionally, incorporating population-level processes into species delimitation may prevent errant findings. A key assumption of the MSCM is that shared genetic polymorphism is a remnant of ancestral polymorphism and not due to gene flow. Indeed, simulation studies have shown that ignoring gene flow leads the MSCM to overestimates of population sizes and underestimates of divergence times (18). For example, Bayesian Phylogenetics and Phylogeography (BPP), a popular implementation of the MSCM for species delimitation (3), has been shown to delimit populations as species even when levels of gene flow between populations are moderate (19, 20), which may not be desirable under certain species concepts (e.g. the Biological Species Concept, 21). It is clear that species delimitation efforts based on the MSCM should proceed with caution, particularly when processes other than lineage divergence are likely to have been important during speciation. However, few studies have considered other parameters when delimiting species (but see 19, 22, which consider migration) primarily due to computational limitations.

Here, we introduce an approach that allows researchers to directly investigate the processes of speciation, ranging from allopatric speciation to founder effect speciation to isolation with secondary contact. We implement this approach in *delimitR*, an R-package that performs demographic model selection under the MSCM. Using a machine-learning algorithm, Random Forest (RF) classification, *delimitR* can evaluate and rank models that include divergence, gene flow, and other demographic processes, such as population size change, any of which may influence the process of lineage splitting that is central to species delimitation. This flexible framework enables researchers to design a model set based on prior knowledge of their focal taxa, and to compare different models of speciation (and models in which a speciation event does not occur). *delimitR* thus allows users to identify the process by which speciation occurred (or did not occur) in their focal taxa. Here, we describe *delimitR*, evaluate its performance using simulations, and apply it to three previously published empirical datasets: the *Hemidactylus* gecko dataset analyzed in Leache *et al*. (23) and two datasets from invertebrates associated with North American pitcher plants analyzed by Satler and Carstens (24).

## Results and Discussion

### Simulation Studies

To evaluate the power of *delimitR*, we designed a simulation study that considered four scenarios in a three-population system (Figure 1a). These scenarios included: 1) no population divergence; 2) divergence between two of the three populations; 3) divergence between all three populations; and 4) divergence between all three populations with secondary contact between the two most closely related populations. We conducted this analysis using both moderate (50,000-100,000 generations) and recent (5,000-10,000 generations) divergence times with 500 to 10,000 unlinked SNPs. The effective number of migrants per generation (Nm) ranged from 0.05 to 5. In the simulations with moderate divergence times, overall error rates were low regardless of the number of SNPs used (0.0009-0.026; Figure 1b). Error rates were zero for the model with no population divergence and highest for the model with three populations and secondary contact (0.0019-0.0672); Figure 1b). In the recent divergence time analyses, overall error rates were moderate regardless of the number of SNPs used (0.13 to 0.18; Table 1; Figure 1b). Error rates were near zero for the model with no population divergence (0 percent) and highest for the three-population model with secondary contact and the three-population model (0.26-0.38; Figure 1b).

**Figure 1:**
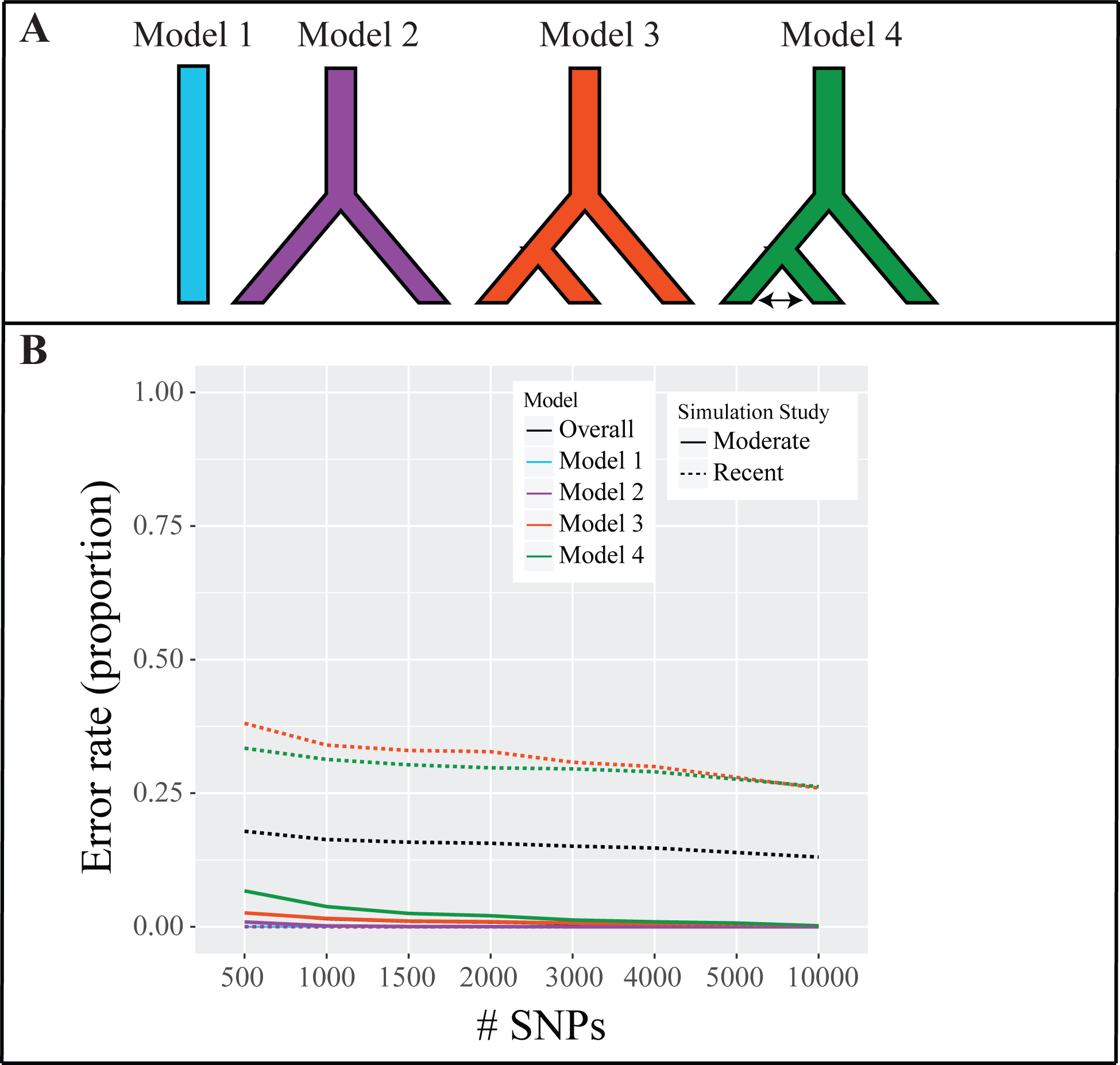
Results of the first simulation study. A) The four models evaluated in the simulation study. B) The results of the simulation study with moderate and recent divergence times. Error rates are reported as the proportion of simulations that were misclassified.

**Table 1:** Error rates from the simulation study for each model and overall. Moderate divergence times were drawn from uniform (50000, 100000) priors. Recent divergence times were drawn from uniform (5000,10000) priors. All error rates in this table are given as proportions of simulations that were misclassified.

| # SNPs | Divergence times | Overall | Model 1 | Model 2 | Model 3 | Model 4 |
| --- | --- | --- | --- | --- | --- | --- |
| 500 | moderate | 0.026 | 0.000 | 0.009 | 0.026 | 0.067 |
| 1000 | moderate | 0.014 | 0.000 | 0.002 | 0.016 | 0.038 |
| 1500 | moderate | 0.009 | 0.000 | 0.001 | 0.011 | 0.025 |
| 2000 | moderate | 0.008 | 0.000 | 0.000 | 0.010 | 0.021 |
| 3000 | moderate | 0.005 | 0.000 | 0.000 | 0.007 | 0.013 |
| 4000 | moderate | 0.004 | 0.000 | 0.000 | 0.005 | 0.009 |
| 5000 | moderate | 0.003 | 0.000 | 0.000 | 0.004 | 0.007 |
| 10000 | moderate | 0.001 | 0.000 | 0.000 | 0.002 | 0.002 |
| 500 | recent | 0.179 | 0.000 | 0.000 | 0.381 | 0.334 |
| 1000 | recent | 0.163 | 0.000 | 0.000 | 0.340 | 0.313 |
| 1500 | recent | 0.158 | 0.000 | 0.000 | 0.330 | 0.303 |
| 2000 | recent | 0.156 | 0.000 | 0.000 | 0.328 | 0.298 |
| 3000 | recent | 0.151 | 0.000 | 0.000 | 0.308 | 0.296 |
| 4000 | recent | 0.147 | 0.000 | 0.000 | 0.300 | 0.290 |
| 5000 | recent | 0.139 | 0.000 | 0.000 | 0.280 | 0.276 |
| 10000 | recent | 0.130 | 0.000 | 0.000 | 0.260 | 0.262 |

Error rates are comparable to those for other species delimitation methods that use genomic data (e.g. BFD* and PHRAPL; 9, 13), and computational times are reasonable. For example, with 10,000 SNPs, four models, and 10,000 simulations per model, the entire analysis carried out in *delmitR* took just over 4 hours of CPU time on six 2.4 GHz processors. The simulated datasets analyzed here are well within the range of those currently being collected by empiricists. Further, even when simulations were conducted with only 500 SNPs (a very low number by current standards), error rates are below 0.05 for a standard species delimitation scenario (Figure 1). These results demonstrate that *delimitR* can achieve comparable or higher power than existing species delimitation methods even while considering more complex demographic scenarios with no loss of computational efficiency.

To evaluate the power of *delimitR* to distinguish among more complex modes of speciation, we used a simulation study based on a continent-island system. Island systems have long been of interest to evolutionary biologists, particularly with respect to speciation. With respect to island systems, founder effect speciation as initially proposed by Mayr (25) involves the founding of an island population by a small number of individuals from a continental source population. Subsequent genetic drift and relaxation of selective constraints lead to a shifts in adaptive peaks, which ultimately result in a speciation event. Though founder effect speciation is one potential driver of divergence in continent-island systems, it is not the only. Other schools of thought hold that vicariance is the main driver of diversification on islands. According to vicariance models, previously contiguous populations are isolated when mainland and island populations separate owing to geological events, and divergence occurs in isolation. Though geographic isolation may seem a key feature of speciation on islands, it has also been proposed that secondary contact could be an important driver of speciation in such scenarios, for example in Darwin’s finches (26). Here we consider six potential scenarios for an island-continental system: 1) no population divergence; 2) allopatric (vicariant) speciation; 3) divergence with secondary contact; 4) divergence with gene flow; 5) founder effect speciation; and 6) founder effects with continuous gene flow (Figure 2a, b). We conducted this analysis using moderate divergence times (50,000-100,000 generations) with the number of unlinked SNPs ranging from 500 to 10,000. Overall error rates were moderate, and decreased when more SNPs were used (0.15-0.32; Table 2; Figure 2c). Error rates were zero for the no divergence model and highest for the founder effect model with gene flow (0.32, 10000 SNPs; Table 2; Figure 2c). Error rates were highest among models that included migration. In particular, error rates were high between the secondary contact model and the founder effect with gene flow model (Figure 1b). This is not surprising, since migration likely swamps the signal of the founder effect, and the patterns predicted by these models converge, making it difficult to distinguish between these scenarios.

**Figure 2:**
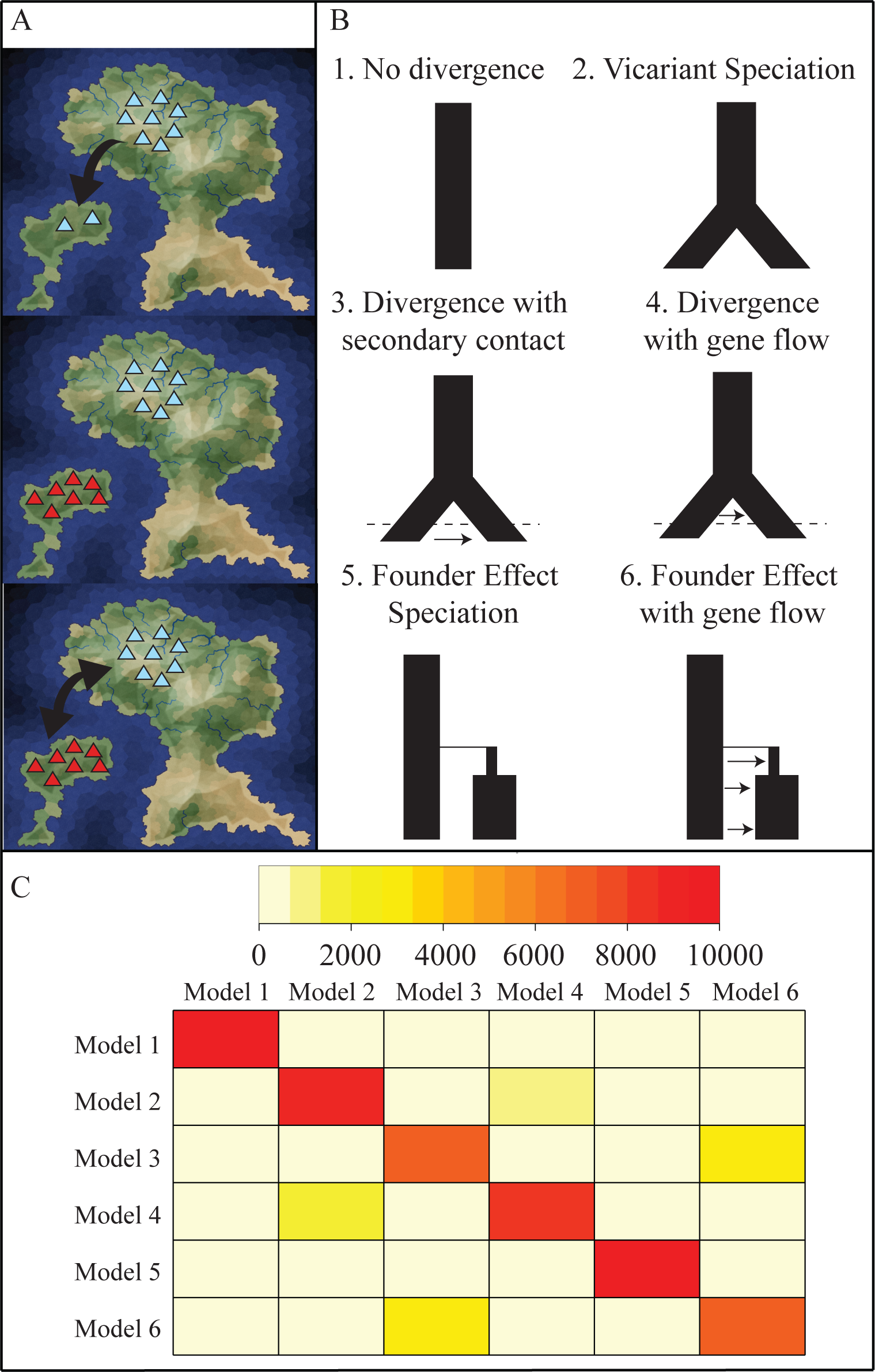
Results of the second simulation study. A) In island-continent systems there are several processes that may contribute to speciation, including: 1) Founder effect speciation, in which a small number of individuals colonizes the island and speciates (top); 2) Vicariant speciation, in which a previously contiguous population diverges into two when island and mainland populations separate (middle); 3) secondary contact, in which a period of initial divergence is followed by a period of gene flow (and potentially reinforcement, or fusion; lower). B) The six models evaluated in the simulation study. C) The heatmap represents the error rates, in terms of the number of simulated datasets that were classified as belonging to a certain model. Each cell represents the number of simulations under the model (row) classified as belonging to each model in the model set (columns). Red along the diagonal indicates that most simulated datasets were correctly classified.

**Table 2:** Error rates from the island-continent simulation study for each model and overall. All error rates in this table are given as proportions of simulations that were misclassified.

| # SNPs | Overall | M 1 | M 2 | M 3 | M 4 | M 5 | M 6 |
| --- | --- | --- | --- | --- | --- | --- | --- |
| 500 | 0.318 | 0.000 | 0.412 | 0.497 | 0.315 | 0.178 | 0.507 |
| 1000 | 0.278 | 0.000 | 0.346 | 0.450 | 0.272 | 0.143 | 0.457 |
| 1500 | 0.260 | 0.000 | 0.315 | 0.436 | 0.253 | 0.130 | 0.423 |
| 2000 | 0.240 | 0.000 | 0.281 | 0.408 | 0.228 | 0.114 | 0.410 |
| 3000 | 0.217 | 0.000 | 0.242 | 0.377 | 0.206 | 0.095 | 0.385 |
| 4000 | 0.205 | 0.000 | 0.210 | 0.355 | 0.200 | 0.085 | 0.379 |
| 5000 | 0.192 | 0.000 | 0.190 | 0.333 | 0.185 | 0.079 | 0.363 |
| 10000 | 0.154 | 0.000 | 0.122 | 0.288 | 0.147 | 0.050 | 0.315 |

### Case Study #1: Hemidactylus geckos

To evaluate the performance of *delimitR* on an empirical dataset, we reanalyzed the data from Leaché *et al*. (23), which consists of five putative species of West African forest geckos *(Hemidactylus fasciatus)*. Previously, Leaché *et al*. found that this dataset consisted of five species using Bayesian Factor Delimitation without considering gene flow, and based on these results, three new species were named. We used priors based on previous results from (23, 27), with migration rates that ranged from 0.05 to 15 Nm. The most recent divergence times were drawn from a uniform (20000,100000) prior distribution. Our results corroborated those of Leaché *et al*. (23), though we considered 13 models, six of which included migration (Figure 3a). The best model was a five population divergence-only model (posterior probability= 0.656). Most models that received votes from the RF classifier included five populations and differed only in whether or not gene flow was included (Figure 3c), and the best two models differed only in one migration parameter (Figure 3b). Overall, we had moderate power to distinguish among the models considered, based on error rates (0.154), with the highest error rates between models that differed only in the inclusion or exclusion of migration parameters (Figure 3c; Table 3). Though error rates are moderate between models that differ only in migration parameters, this is not surprising given that migration rates as low as 0.05 Nm were considered in this analysis.

**Figure 3:**
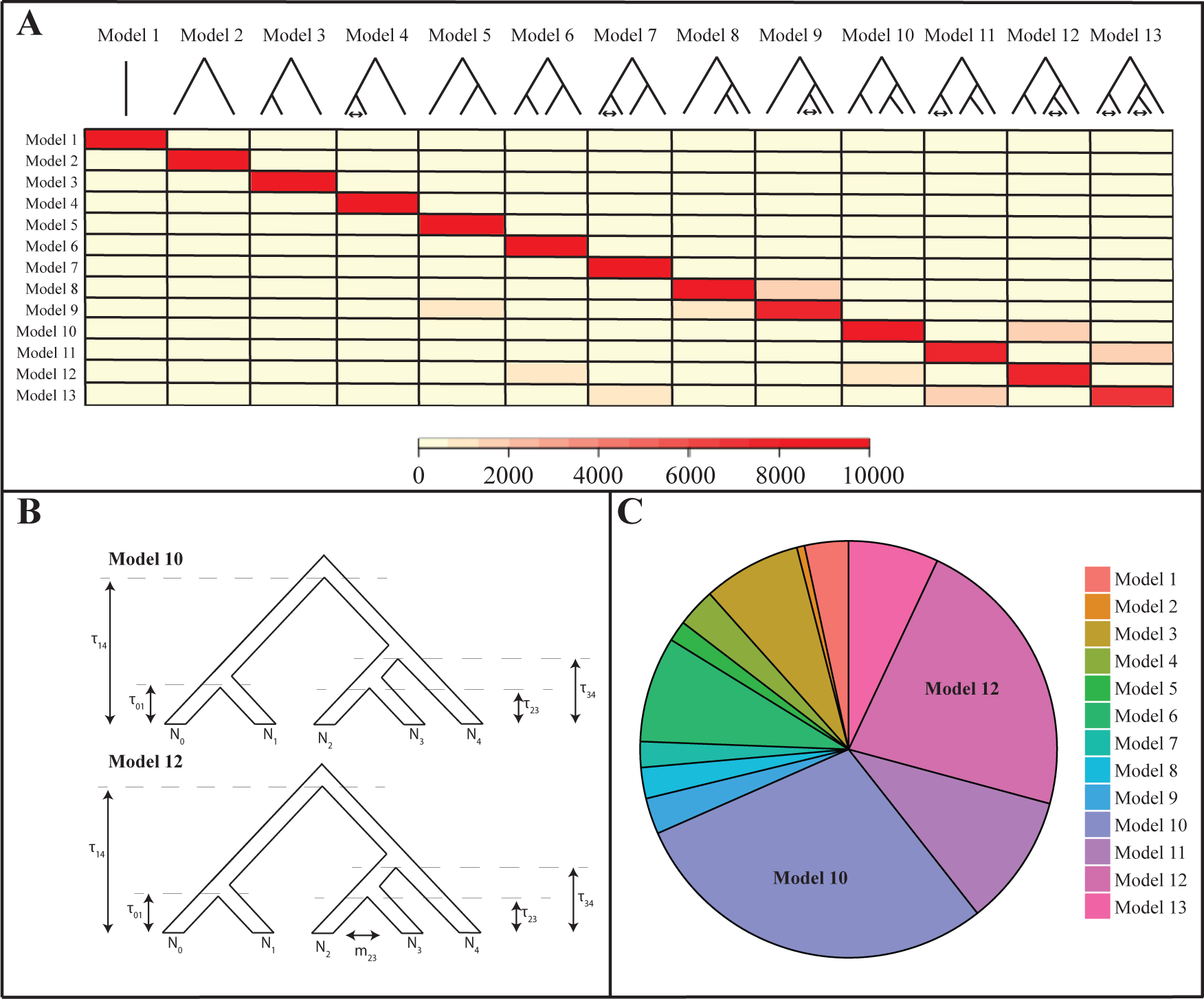
Results from the analysis of the *Hemidactylus* geckos dataset. A) The thirteen models evaluated in the study. The heatmap represents the error rates, in terms of the number of simulated datasets that were classified as belonging to a certain model. Each cell represents the number of simulations under the model (row) classified as belonging to each model in the model set (columns). Red along the diagonal indicates that most simulated datasets were correctly classified. B) The two best models. The populations (0–5) correspond to *H. fasciatus, H. kyaboboensis, H. coalescens*, Bioko Island, and *H. eniangii*, respectively. N0 represents the population size of population 0 *(H. fasciatus)*. τ_01_ represents the divergence time (in number of generations) between population 0 (*H. fasciatus*) and population 1 (*H. kyaboboensis*), and m01 represents migration between populations 0 and 1. C) The proportion of decision trees in the RF classifier that voted for each model when the classifier was applied to the observed data.

**Table 3:**
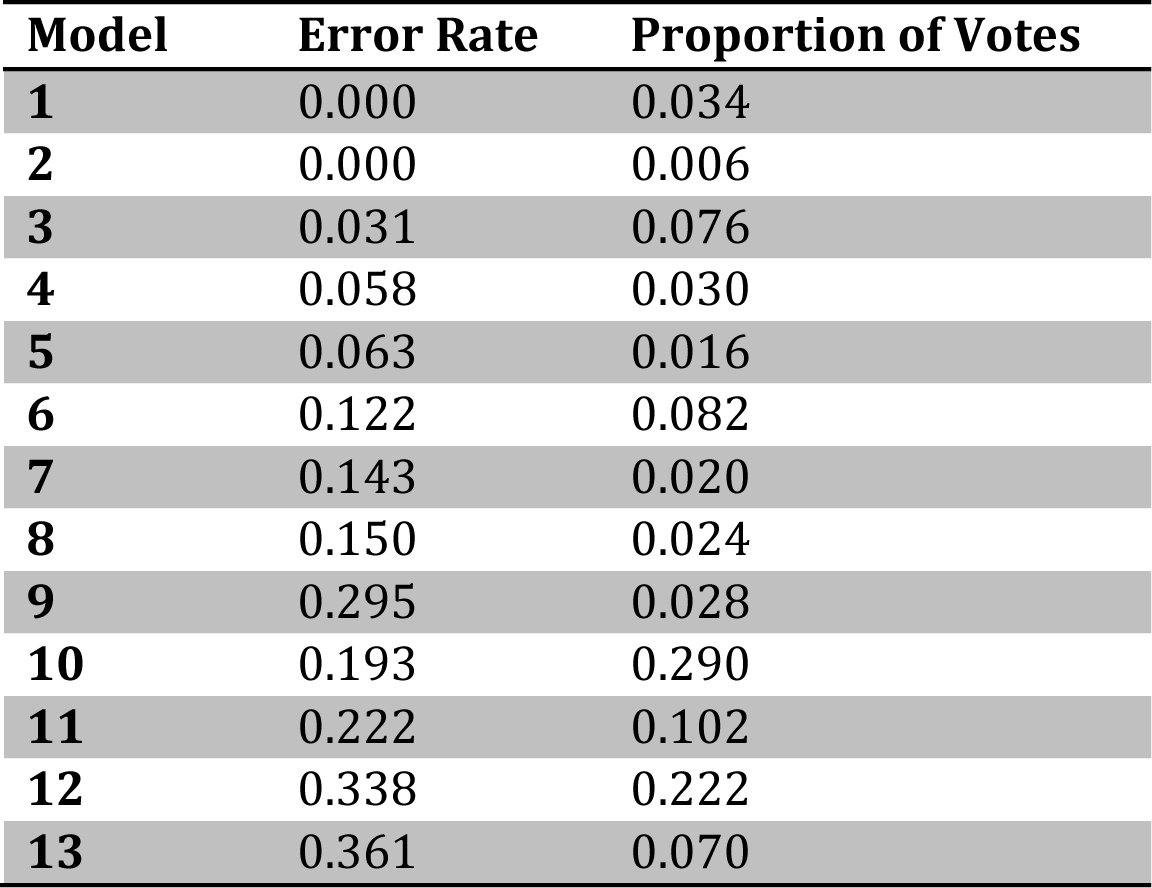
Results from the *Hemidactylus* geckos. Model numbers correspond to those reported in Figure 2. All error rates in this table are given as proportions of simulations that were misclassified. The proportion of trees in the RF classifier that voted for each model when the classifier was applied to the observed data.

| **Model** | **Error Rate** | **Proportion of Votes** |
| --- | --- | --- |
| **1** | 0.000 | 0.034 |
| **2** | 0.000 | 0.006 |
| **3** | 0.031 | 0.076 |
| **4** | 0.058 | 0.030 |
| **5** | 0.063 | 0.016 |
| **6** | 0.122 | 0.082 |
| **7** | 0.143 | 0.020 |
| **8** | 0.150 | 0.024 |
| **9** | 0.295 | 0.028 |
| **10** | 0.193 | 0.290 |
| **11** | 0.222 | 0.102 |
| **12** | 0.338 | 0.222 |
| **13** | 0.361 | 0.070 |

Another benefit of *delimitR* is the ease with which uncertainty is quantified. When model selection is performed in *delimitR*, a table with estimated error rates is automatically generated. These error rates can help the user understand how much power they have to distinguish among similar models without necessitating a full power analysis. Here, it is clear that error rates are moderate between models differing in migration parameters (Figure 3a). This encourages caution in interpreting the results of model selection, and would encourage the researcher to use a secondary method (e.g. fastsimcoal2; 11) to quantify migration rates under the best divergence model and evaluate whether or not the magnitude of migration was biologically meaningful.

### Case Study #2: Sarracenia pitcher plant invertebrates

To further evaluate the performance of *delimitR*, we reanalyzed two datasets from Satler and Carstens (24). The datasets are from two ecological associates of North American pitcher plants: a moth *(E. simicrocea)* and a spider *(P. viridans). E. simicrocea* is an obligate inquiline commensal with the pitcher plant *S. alata*, while *P. viridans* is an opportunistic capture interrupter (24). Both datasets consisted of two potential species for each nominal species: one east of the Mississippi River and another west of the Mississippi River. Previous results suggest high migration rates between populations east and west of the Mississippi for the spider *P. viridans* and low migration rates between populations east and west of the Mississippi for the moth *E. semicrocea*. We considered four models: a one-population model, a two-population divergence only model, a two population model with secondary contact, and a two population model with migration during the initial stages of divergence (Figure 4). For both species, *delimitR* supports migration models, and has high power to distinguish among models (0.045 and 0.050 error rates for *E. semicrocea* and *P. viridans*, respectively; Figure 4, Table 4). For *P. viridans*, the best model includes migration following secondary contact (pp=0.944), while for *E. semicrocea*, the best model includes migration during the initial stages of divergence (pp=0.798) (Figure 4). For *E. semicrocea*, the error rates generated by *delimitR* indicate that it is sometimes difficult (0.073-0.093 error rate) to distinguish among the secondary contact and divergence with gene flow models, encouraging caution in interpreting the results regarding the timing of migration. For *P. viridians*, error rates are low except for the model of divergence with gene flow, which is mistaken for a two-population model 12 percent of the time. Given that we did not select either of these models, and that error rates for the selected model are low (0.0009), we have high confidence in our results for this species. The results from these empirical studies highlight that results from *delimitR* are largely concordant with those from other popular programs, and can lend further insight into the processes driving speciation.

**Figure 4:**
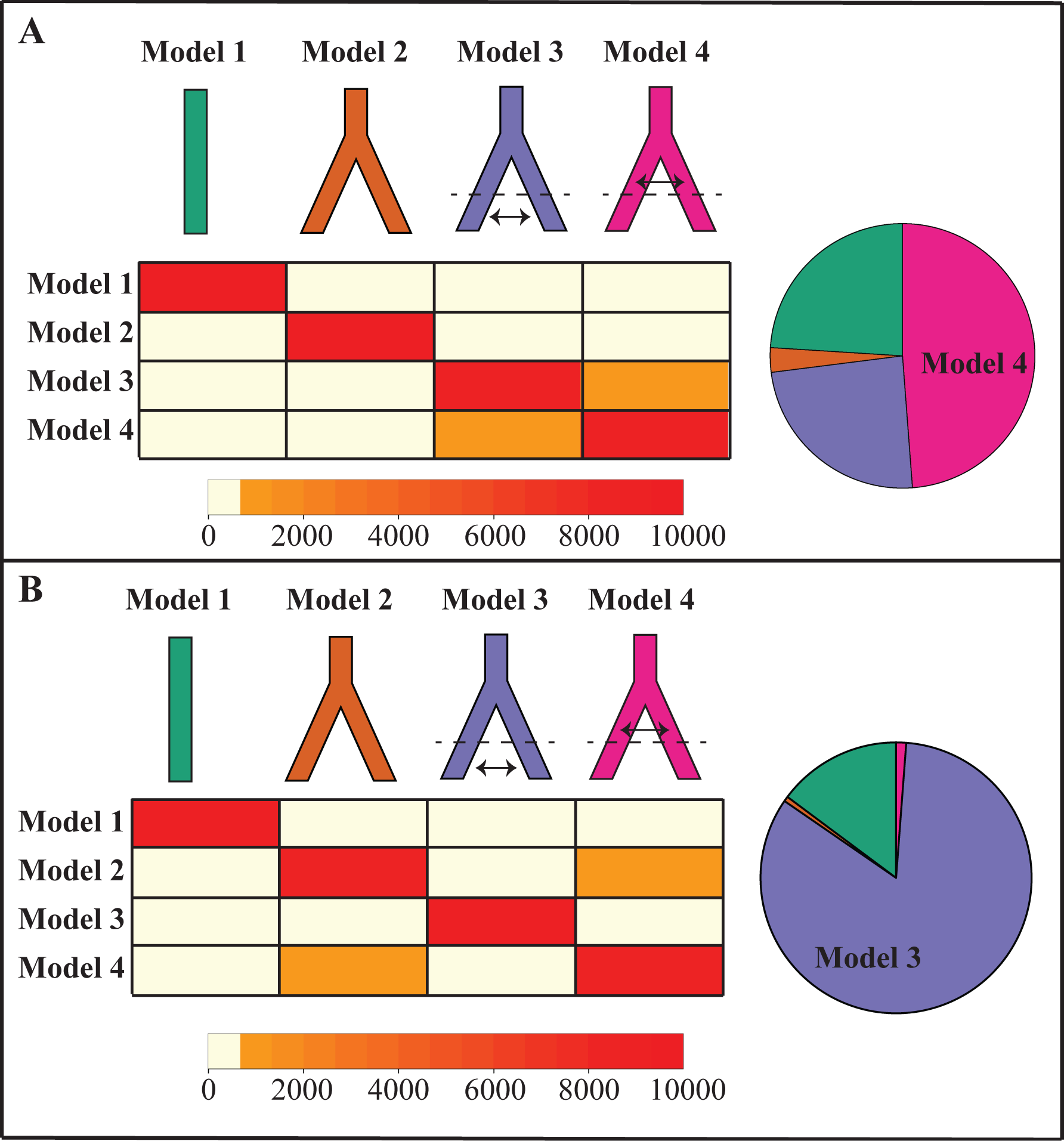
Results from the analysis of the two pitcher plant invertebrates. The four models are drawn. The heatmap represents the out of the bag error rates, in terms of the number of simulated datasets that were classified as belonging to a certain model. Each cell represents the number of simulations under the model (row) classified as belonging to each model in the model set (columns). Red along the diagonal indicates that most simulated datasets were correctly classified. The pie charts indicate the proportion of decision trees in the RF classifier that voted for each model when the classifier was applied to the observed data. A) Results from *E. semicrocea* and B) results from *P. viridans*.

**Table 4:** Results from the two pitcher plant-associated invertebrates. Model numbers correspond to those reported in Figure 3. All error rates in this table are given as proportions of simulations that were misclassified. The proportion of trees in the RF classifier that voted for each model when the classifier was applied to the observed data.

| Species | Model | Error rates | Proportion of votes |
| --- | --- | --- | --- |
| *E. semicrocea* | 1<sup>1</sup> | 0.0000<sup>2</sup> | 0.240<sup>2</sup> |
| *E. semicrocea* | 2 | 0.0141 | 0.030 |
| *E. semicrocea* | 3 | 0.0930 | 0.242 |
| *E. semicrocea* | 4 | 0.0735 | 0.488 |
| *P. viridans* | 1 | 0.0002 | 0.148 |
| *P. viridans* | 2 | 0.0802 | 0.006 |
| *P. viridans* | 3 | 0.0009 | 0.834 |
| *P. viridans* | 4 | 0.1175 | 0.012 |

## Conclusions

### Understanding the Speciation Process

In comparison to previous approaches to species delimitation, *delimitR* allows unmatched flexibility by enabling users to focus on the process by which species have formed. Notably, *delimitR* can be applied to evaluate demographic models consistent with a variety of types of speciation under nearly any species concept. Researchers can use their expertise and familiarity with the focal system to identify reasonable priors on divergence times and migration rates, as well as to decide which models should be included in the model set. After model selection is complete, researchers can use the selected model to estimate parameters, and this information can be used to assess whether or not the divergences in the model represent true speciation events. A secondary benefit of *delimitR* is that it will require researchers to be clear and explicit about what constitutes a species, and thereby should increase transparency and repeatability in species delimitation studies by connecting the delimited species to the process by which they formed.

### Machine Learning and Genomic data

Computational limits are perhaps the primary reason that previous approaches to species delimitation have not focused on the process of speciation. To circumvent this issue, *delimitR* uses a machine-learning algorithm (RF). The use of the RF classifier and the binned multidimensional folded site frequency spectrum (mSFS) allow us to compare a large set of models using large datasets in hours of computational time, a task unmanageable using standard approaches to model comparison. Furthermore, the RF approach used here automatically generates estimated error rates with almost no additional computational expense, giving researchers a built in way to assess power, without needing to perform a full power analysis. This will encourage a nuanced interpretation of results when the power to choose among the models in the model set is low, and will prevent researchers from blindly comparing models among which they do not have enough data to distinguish. Due to the use of the RF classifier, this method is applicable to datasets with moderate number of populations (up to 5 evaluated here), moderate-to-large numbers of SNPs (up to 10,000 evaluated here), and moderate numbers of individuals per population, regardless of whether or not researchers have access to high-power computing resources.

## Materials and Methods

### Generating the Model Set

By default, *delimitR* considers models of divergence and divergence with secondary contact. It requires that the user provide a guide tree (or a list of guide trees) and a maximum coalescence interval (max coal) in which to consider gene flow prior to the generation of the model set. The max coal parameter dictates how far into the past secondary contact will be considered (e.g., if max coal is set to one, then *delimitR* will only generate models that consider gene flow in the present). Additionally, the user must provide information on divergence time priors, population size priors, and migration rate priors. Given this information, *delimitR* will produce fastsimcoal2 (fsc2; 27) input files for the default model set with the function setup_fsc2(). For example, if there were three populations, the user provided guide tree indicating that populations 0 and 1 were more closely related to each other than to population 2, and the max coal parameter was set to one, then the model set generated would consist of four models: a single population model, a two population model, a three population model, and a three population model with secondary contact between populations 0 and 1 (Figure 1a). Though the default model set may be limited in some applications, it is straightforward for the user to include models outside of this set. The user need only generate the fsc2 input files by hand and place them in the working directory. This flexibility makes *delimitR* applicable to any range of demographic scenarios that can be implemented in fsc2.

### Simulating and summarizing data

We use fsc2 (28), a coalescent simulator, to simulate data under each model in the model set. We simulate the mSFS using parameters drawn from user-specified priors. We do not consider the monomorphic cell when simulating mSFS, and we use only unlinked SNPs. Given a user-specified number of replicates *N, delimitR* will simulate *N* replicates under each model in the model set and output one mSFS per simulation using the function fastsimcoalsims().

When all polymorphic sites are unlinked and biallelic (29), the mSFS is a complete summary of the data. However, inferences based on the mSFS may be inaccurate when too few segregating sites are sampled (30). To address this problem, we used a binning strategy to further summarize the mSFS following Smith *et al*. (31) to generate the binned SFS (bSFS). We apply the same binning strategy to the simulated and observed data using the functions makeprior() and prepobserved(), respectively.

### Building the Random Forest classifier and assessing power

We used the simulated data to construct a Random Forest (RF) classifier, in which the bins of the bSFS are the predictor variables and the model used to simulate the data is the response variable. To build the RF classifier, *delimitR* uses the R package “abcrf’ (32). The RF classifier consists of a user-defined number *N* of decision trees. Each decision tree is constructed from a subset of the prior, and at each node in each decision tree a bin of the bSFS is considered, and a binary decision rule is constructed based on the number of SNPs in the bin. When this classifier is applied to new datasets, we move down nodes until we reach the leaves of the tree, which, in this case, are model indices. Each decision tree votes for a model, and the model receiving the largest portion of votes is selected as the best model.

We use out-of-the-bag (oob) error rates to assess the power of the RF classifier. Since only a portion of the prior is used for the construction of each decision tree, we can take an element of the prior, consider only decision trees constructed without reference to that element, and calculate how often we choose an incorrect model. The construction of the RF classifier and calculation of oob error rates is implemented in *delimitR* with the function RF_build_abcrf().

### Classifying the observed data

We use the predict.abcrf() function from the R package “abcrf’ (32), as implemented by the RF_predict_abcrf() function in *delimitR* to select the model that simulates data most similar to the observed data. We also estimate the posterior probability of the selected model by regressing against oob error rates following Pudlo *et al*. (32). Notably, *delimitR* is applicable to a wide range of datasets and systems. Any data that can be used to build a folded multidimensional SFS can be used in *delimitR;* this includes data from organisms with various levels of ploidy and datasets with no outgroup sequence available.

### Simulation study

To assess the accuracy of our method, we performed two simulation studies. All analyses were carried out on the Ohio Super Computer (33). For the first simulation study, we focused on a model with three species, and considered migration during the first coalescent interval, resulting in four models under the default model set generated by *delimitR* (Figure 1a). We sampled 10 diploid individuals from each population (20 alleles). Population sizes were drawn from uniform (10000,100000) priors. Divergence times between species 0 and 1 were drawn from a uniform (50000,100000 generations) prior for the moderate divergence time study and from a uniform (5000, 10000 generations) prior for the recent divergence time study, and divergence times between the ancestor of species 0 and 1 and species 2 were drawn from a uniform (500000, 1000000 generations) prior for the moderate divergence times study and from a uniform (50000, 100000 generations) prior for the recent divergence times study. The migration rates were drawn from a uniform (0.000005, 0.00005) prior, corresponding to 0.05 to 5 Nm. We simulated 10,000 datasets under each of the four models for both the moderate and recent divergence time analyses. We evaluated the performance of the RF classifier using datasets containing 500, 1000, 1500, 2000, 3000, 4000, 5000, and 10000 SNPs. We used ten classes per population to summarize the mSFS. We constructed an RF classifier and calculated oob error rates in *delimitR*. In addition to oob error rates, we used a cross-validation approach to assess the accuracy of our classifier. We simulated 1,000 pseudoobserved datasetes under each of the four models. We applied the RF classifier constructed above to each of these datasets, and calculated how often the correct model was selected for each dataset (Supporting Information: Tables S1 and S2).

The second simulation study focused on a continent-island system, and considered six models (Figure 2b). We sampled 10 diploid individuals from each population (20 alleles). Population sizes were drawn from a uniform (50000, 100000) prior for the island population and a uniform (75000, 20000) prior for the continental population. For models that included divergence, divergence times between the island and continental populations were drawn from a uniform (50000, 100000 generations) prior. For models with migration, the migration rate was drawn from a uniform prior that corresponded to 0.05 to 10 Nm. For models including a bottleneck, the proportion of the population that remained during the bottleneck was drawn from a uniform (0.001, 0.01) prior, which corresponds to 50 to 100 individuals, and the bottleneck lasted from 100 to 500 generations. We evaluated the performance of the RF classifier using datasets containing 500, 1000, 1500, 2000, 3000, 4000, 5000, and 10000 SNPs. We used ten classes per population to summarize the mSFS. We constructed an RF classifier and calculated oob error rates in *delimitR*. In addition to oob error rates, we used a cross-validation approach to assess the accuracy of our classifier. We simulated 1,000 pseudoobserved datasetes under each of the six models. We applied the RF classifier constructed above to each of these datasets, and calculated how often the correct model was selected and estimated the posterior probability of the selected model for each dataset (Supporting Information: Table S3).

### Case Study #1: Hemidactylus geckos

To assess the power and accuracy of *delimitR* when applied to real data, we reanalyzed the data from Leache *et al*. (23). A custom python script was used to construct the mSFS from the BEAST input file (available on github). We used a threshold of 50%, meaning only SNPs that occurred in at least 50% of individuals in each population were used to build the SFS. When SNPs were present in more than 50% of individuals, we randomly sampled alleles from each population. We excluded one individual from this dataset (ENG_CA2_20), as it is thought to have been mislabeled (A. Leache, pers. communication). We used the folded SFS, which does not require information about whether or not an allele is derived or ancestral, and used ten classes per population to summarize the mSFS.

We considered all collapsed histories and histories with migration parameters in the first coalescent interval (between *H. fasciatus* and *H. kyaboboensis* and between *H. coalescens* and Bioko Island). This resulted in a total of 13 models (Figure 3a). As a guide tree, we used the tree published in (23). Priors were chosen based on the results from (23, 27). All priors were drawn from uniform distributions. For *H. fasciatus, H. kyaboboensis*, and *H. coalescens*, population size priors were converted from the coalescent units reported in Leache and Fujita, using a mutation rate of 5.29E-9 (mean mtDNA rate for reptiles; 20) and a generation time of one year. These values were not estimated for *H. eniangii* and the Bioko Island populations separately in (27). For these two populations, we used broad uninformative priors. Divergence times were converted from coalescent units using the same mutation rate as above. The divergence time between *H. coalescens* and the Bioko Island population was not estimated by (27) so a broad uninformative prior compatible with the rest of the tree was used. Migration rates were drawn from a uniform (0.000001, 0.00005) prior. More information on the priors is given in Supporting Table S1. We simulated 10,000 mSFS under each model. We then summarized the simulated and observed mSFS by binning (five classes per population), built an RF classifier, and applied this classifier to the observed data using functions in the R package *delimitR* as described above.

### Case Study #2: Sarracenia pitcher plant invertebrates

Finally, we applied the method to two datasets from Satler and Carstens (24): a moth and a spider. A custom python script was used to construct the mSFS from the pyrad output files (available on github). We used a threshold of 50%, meaning only SNPs that occurred in at least 50% of individuals in each population were used to build the SFS. When SNPs were present in more than 50% of individuals, we randomly sampled alleles from each population. We used the folded SFS, which does not require information about whether or not an allele is derived or ancestral. We used ten classes per population to summarize the SFS.

For both the moth and the spider, we considered two potential species: one east of the Mississippi and one west of the Mississippi. To generate priors for divergence times, migration rates, and population sizes, we doubled the width of the confidence intervals reported in Satler and Carstens (15; Table 3). We considered four models: a one population model, a two population model, a two population model with secondary contact, and a two population model in which gene flow occurred during divergence. We simulated 10,000 mSFS under each model, summarized the simulated and observed mSFS by binning, built an RF classifier, and applied this classifier to the observed data using functions in the R package *delimitR*.

### Data Availability

The R-package *delimitR*, as well as a full tutorial is available on github (https://github.com/meganlsmith/delimitR).

## Acknowledgements

MLS was funded by a NSF GRFP (DGE-1343012), and this work was funded by NSF (DEB1457519). We would like to thank members of the Carstens lab, Ryan Garrick, and Benjamin Stone for comments that improved the manuscript prior to publication, and we would like to thank Paul Blischak for the name *delimitR*. We would like to thank the Ohio Supercomputer for computing resources (allocation grant PAS1181-2).

## Supporting Information

**Supporting Table S1:** Cross-validation results for simulation study #1 with moderate divergence times with 10,000 SNPs.. The numbers are reported in each row are the number of datasets simulated under the general model that were classified as belonging to each of the models.

| **Generating Model** | **Model 1** | **Model 2** | **Model 3** | **Model 4** |
| --- | --- | --- | --- | --- |
| **Model 1** | 1000 | 0 | 0 | 0 |
| **Model 2** | 0 | 1000 | 0 | 0 |
| **Model 3** | 0 | 0 | 998 | 2 |
| **Model 4** | 0 | 0 | 3 | 997 |

**Supporting Table S2:** Cross-validation results for simulation study #1 with recent divergence times with 10,000 SNPs.. The numbers are reported in each row are the number of datasets simulated under the general model that were classified as belonging to each of the models.

| **Generating Model** | **Model 1** | **Model 2** | **Model 3** | **Model 4** |
| --- | --- | --- | --- | --- |
| **Model 1** | 1000 | 0 | 0 | 0 |
| **Model 2** | 0 | 1000 | 0 | 0 |
| **Model 3** | 0 | 0 | 732 | 268 |
| **Model 4** | 0 | 0 | 243 | 757 |

**Supporting Table S3:**
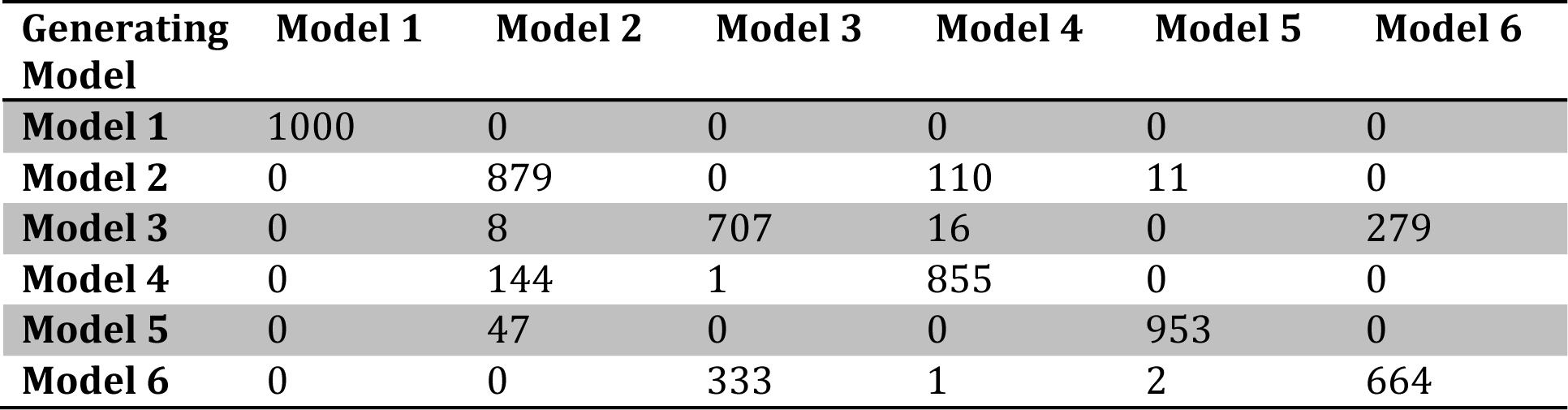
Cross-validation results for the island-continent simulation study with 10,000 SNPs̤ The numbers are reported in each row are the number of datasets simulated under the general model that were classified as belonging to each of the models.

| **Generating Model** | **Model 1** | **Model 2** | **Model 3** | **Model 4** | **Model 5** | **Model 6** |
| --- | --- | --- | --- | --- | --- | --- |
| **Model 1** | 1000 | 0 | 0 | 0 | 0 | 0 |
| **Model 2** | 0 | 879 | 0 | 110 | 11 | 0 |
| **Model 3** | 0 | 8 | 707 | 16 | 0 | 279 |
| **Model 4** | 0 | 144 | 1 | 855 | 0 | 0 |
| **Model 5** | 0 | 47 | 0 | 0 | 953 | 0 |
| **Model 6** | 0 | 0 | 333 | 1 | 2 | 664 |

**Supporting Table S4:** Hemidactylus priors. coal/bioko indicates the common ancestor of coal and bioko. coal-bioko indicates the divergence event between coal and bioko. Population sizes are given in terms of the number of haploid individuals. Divergence times are given in generations before the present.

| **Population Size Priors** |  |
| --- | --- |
| **Population** | **Prior** |
| **fas** | (50000, 100000) |
| **kya** | (50000, 100000) |
| **coal** | (250000, 300000) |
| **Bioko** | (50000, 300000) |
| **Enj** | (50000, 300000) |
| **Divergence Time Priors** |  |
| fas-kya | (20000, 50000) |
| coal-bioko | (20000, 50000) |
| coal/bioko-eni | (100000, 200000) |
| coal/bioko/eni-fas/kya | (300000, 400000) |

